# Inferring cell state by quantitative motility analysis reveals a dynamic state system and broken detailed balance

**DOI:** 10.1101/168534

**Authors:** Jacob C. Kimmel, Amy Y. Chang, Andrew S. Brack, Wallace F. Marshall

## Abstract

Cell populations display heterogeneous phenotypic states at multiple scales. Similar to molecular features commonly used to explore cell heterogeneity, cell behavior is a rich phenotypic space that may allow for identification of relevant cell states. Inference of cell state from cell behavior across a time course may enable the investigation of dynamics of transitions between heterogeneous cell states, a task difficult to perform with destructive molecular observations. Cell motility is one such easily observed cell behavior with known biomedical relevance. To investigate cell heterogeneity through the lens of cell behavior, we developed Heteromotility, a software tool to extract quantitative motility features from timelapse cell images. In mouse embryonic fibroblasts (MEFs), myoblasts, and muscle stem cells (MuSCs), Heteromotility analysis identifies multiple motility phenotypes within the population. In all three systems, the motility state identity of individual cells is dynamic. Quantification of state transitions reveals that MuSCs undergoing activation transition through progressive motility states toward the myoblast phenotype. By probability flux analysis, we find that this MuSC motility state system breaks detailed balance, while the MEF and myoblast systems do not. Our data indicate that the system regulating cell behavior can be decomposed into a set of attractor states which depend on the identity of the cell, together with a set of transitions between states governed by inputs from signaling pathways such as oncogenes and growth factors. Within one state, equilibrium formalisms can capture variation in behavior, while switching between states violates equilibrium conditions and would require an external driving force. These results support a conceptual view of the cell as a non-deterministic state automaton, responding to inputs from signaling pathways and generating outputs in the form of observable motile behaviors.

## Introduction

Within any group of cells, each individual cell is not necessarily like its neighbors. These differences often have functional significance, with cells at different points along the phenotypic spectrum exhibiting distinct behavior [1]. This has been noted across a broad swath of cell biology, including in the cases of stem cell biology [2,3] and cell geometry definition [4]. Regulatory decisions at one scale may reflect phenotypes at other scales, allowing identification of a broad “cellular state” based on a more limited set of observations. Effective classification of cancerous cell functionality based on morphology demonstrates this concept [5-7]. Since cell state determines cell function, state transitions are manifest as changes in cell behavior. Understanding the regulation of cell behavior will require understanding the nature of the cell state-space and the transitions that take place within it. Is the state space continuous or discrete? Are all state transitions equally likely, as in an equilibrium system, or do they tend to take place in a specific sequence, as in computing device? Answering such questions requires a framework for defining cell state in terms of observable behaviors. Regardless of the method used to probe cell state, it must be able to measure state in living cells at multiple time points, in order to allow state transitions to be characterized.

Recent advances in single cell assays have allowed for detailed, quantitative descriptions of individual cells at the molecular level. Single cell sequencing technologies in particular have uncovered heterogeneity at both the transcriptional and epigenetic level [8,9]. In multiple stem cell compartments, molecular analysis of heterogeneity has revealed that not all stem cells are functionally equivalent. Within the hematopoietic [10], muscle [11], epithelial, and other [12] stem cell pools, subpopulations of cells with different functionality coexist. Stem cell heterogeneity has been demonstrated in the form of lineage-bias or differences in regenerative capacity. A similar phenomena is present in malignant tumors. Within a given tumor, some subpopulations may have higher tumorogenic potential, or increased resistance to a particular therapy [13,14]. Understanding heterogeneity in these and other contexts is essential to building an accurate picture of cell-based systems.

In addition to defining cell states to account for population heterogeneity, we also seek to understand the transitions between states, because those transitions reveal the logic of the cellular control system. In seminal work, Waddington introduced the conceptual model of an ‘epigenetic landscape’ governing cell phenotype decisions, akin to a potential energy landscape in a physical system [15,16]. In this model, cells progress through phenotypic states by migrating continuously down the gradient of the landscape, eventually resting in a stable basin of attraction. This model implies that cell state transitions are governed by a ‘potential energy’ in each state, which can be estimated by the state’s stability. A cell system governed by this model would display detailed balance in the absence of an external force or signal, and break detailed balance in the presence of such an external input [16,17]. By directly measuring the dynamics of cell state transitions, we can produce an estimate of the landscape for cell behavioral phenotypes and determine if state transitions occur stochastically or are influenced by external inputs.

While existing molecular assays such as single-cell RNA-sequencing can provide detailed information about a cell population’s heterogeneity, these assays are generally destructive and restricted to a single time point for analysis. Methods have been developed to infer cell state lineages from observations at a single time point[18,19], but these methods are not able to quantify dynamics and assume that proximity in the measured variable space describes a transition relationship between states. Real-time, non-destructive assays that reveal subpopulation composition over time and observe state transitions in living cells would serve as complementary approaches to investigate cellular states and state transitions.

Cells in culture have a diverse behavioral repertoire. Each cell may exhibit motion, dynamic morphology changes [4,20], and symmetric or asymmetric divisions, all of which can be observed by simple microscopy. These cell behaviors represents a rich phenotypic space from which many quantitative features may be extracted. Behavior also has an inherent functional relevance at the single cell level, and often a functional relevance on the system level. For example, extracellular matrix remodeling and deposition by fibroblasts is dictated by their motion [21,22], and stem cell and progenitor migration is critical for organismal development [23]. Within each cell, behavior represents a layer of abstraction above molecular phenotypes. In a sense, the “sum” of a cell’s molecular organization at each level produces these observable behaviors. Hence, observably different behavioral states may serve as a proxy for distinct ensembles of underlying molecular states.

Cell motility is particularly dramatic and yet difficult to examine quantitatively. Traditional cell motility assays rely upon a binary filtration of cells based on a functional test, such as crossing a membrane barrier. Timelapse microscopy has also been appreciated for decades as a means of tracking and quantifying cell motility at the single cell level [24-27]. These classic studies demonstrate that cell motility is predictive of cellular function and that quantitative motility analysis can elucidate underlying cell state control mechanisms [26,27]. Recent approaches to timelapse motility analysis have expanded upon these techniques to extract multidimensional quantitative information from individual cells [28-33]. However, existing methods focus largely on speed and distance metrics of cell motility on a single arbitrarily-chosen timescale, limiting the degree of heterogeneity that can be revealed within a population. Here, we present Heteromotility, a software tool for quantitative analysis of cell motility in timelapse images with a diverse feature set. In addition to commonly calculated features such as distance traveled, turning, and speed metrics, Heteromotility provides features that allow for comparisons to models of complex motion, such as Levy flights and fractal Brownian motion, and estimate long-term dependence within a cell’s displacement distribution. This feature set creates a high-dimensional space representing the possible phenotypes of cell motility. We provide tools to map this high-dimensional feature space into a low dimensional state space to quantify changes in cell motility phenotypes over time as cell state transitions. These tools allow the dynamics of cell behavior states to be elucidated, in addition to simply identifying heterogenous phenotypes.

We demonstrate that Heteromotility analysis is sufficient to discriminate between simulations of several models of motion. Applied to wild-type and transformed mouse embryonic fibroblasts (MEFs), Heteromotility analysis reveals a shared set of motility states between the two systems, in which transformed cells preferentially occupy a more motile state. In mouse myoblasts and MuSCs, phenotypically distinguished motility states are revealed. MuSCs display a series of progressively more motile states, suggesting states of activation. Quantifying transition dynamics within this state system over time reveals that MuSCs transition through state space toward a progressively more activated, myoblast-like motility phenotype. Viewed through the lens of myogenic activation, we are able to follow the activation dynamics of individual MuSCs for the first time. Applying probability flux analysis, we find that the MuSC motility state system breaks detailed balance, quantitatively confirming that these state transitions occur in an ordered and predictable sequence, as in a computing device.

## Results

### Heteromotility analysis approach

Heteromotility analyzes motility features in a set of provided motion paths, as obtained through timelapse imaging, image segmentation, and tracking (see Methods). From these paths, 79 motility features are calculated to comprise a “motility fingerprint” (Fig. S1). These features include simple metrics of speed, total and net distance traveled, the proportion of time a cell spends moving, and the speed characteristics during that period. The linearity of motion is assessed by linear regression through all points occupied by the cell in the time series, taking Pearson’s *r*^2^ as a metric of fit. Monotonicity is also considered for the distribution of points, using Spearman’s *ρ*^2^. In some instances, cells have been proposed to have a directional bias when making turns [34]. Turn direction and magnitude features are provided that consider turns on various time intervals.

Another class of features is concerned with the directionality and persistence of motion. There are several possible ways to characterize the persistence of motion, hence we have included several metrics that allow for reasonable consideration. A cell’s “progressivity,” is considered as the ratio of net distance to total distance traveled. This serves as a metric of directional persistence [35]. Mean squared displacement (MSD) is considered for each cell for a variable time lag *τ*, and the power law exponent *α* is taken as a quantification of the relationship *MSD* ∝ *τ^α^*. Directed motion may therefore be expected to have a larger value than undirected random-walk motion or non-motion, indicating a superdiffusive behavior [36].

The distribution of a cell’s displacement steps is also informative. Motion that exhibits a displacement distribution with a heavy right tail, referred to as a Levy flight, has been demonstrated to optimize the success of a random search [37]. This property has led to the Levy flight foraging hypothesis, suggesting that biological systems may perform Levy flight-like motion when searching for resources [38-40]. To assess the Levy flight-like nature of a cell’s motility behavior, the Heteromotility software provides metrics of displacement kurtosis for displacement distributions on multiple time scales. Larger values of kurtosis indicate heavier distribution tails, such that higher kurtosis may indicate more Levy flight-like motion. The Heteromotility software also considers the non-Gaussian parameter α_2_, for which larger values indicate a more heavily tailed, Levy flight-like distribution [41,42]. The second and third moments of the displacement distribution are also provided as features.

If displacements are considered as a time series, the self-similarity and long range memory may provide insight into the coordination of motility behavior [26]. The Heteromotility software calculates the autocorrelation function for displacements with variable time lags *τ* as a metric of self-similarity. Fractal Brownian motion (fBm) describes a Gaussian process *B_H_*(*t*) for a time *t* on the continuous interval [0, *T*] with successive displacements that are not necessarily independent. The Hurst parameter *H* describes the self-similarity of a fBm process, with the interval 0 < *H* < 0.5 describing a process with negatively correlated successive displacements (large displacements are more likely to follow small displacements), *H* = 0.5 describing non-correlated, independent successive displacements (Brownian motion), and 0.5 < *H* < 1 describing positively correlated successive displacements (large displacements are more likely to follow large displacements)(see Supp. Methods for additional detail). The Heteromotility software estimates the Hurst parameter as a metric of long range memory using Mandelbrot’s rescaled range method (see Supp. Methods) [43,44]. As Brownian motion displays a Hurst parameter *H* = 0.5, deviations from this value may indicate long range memory of a cell’s displacement series and coordinated motility behavior.

### Heteromotility features are sufficient to distinguish canonical models of motion

In principle one could devise an infinite number of motion descriptors. How can we determine whether a given set of motion features is sufficient to capture meaningful aspects of motion? To test whether the feature set outlined above is sufficient to distinguish biologically relevant models of motion, we simulated and analyzed cell paths generated by four distinct motion models: unbiased random walks, biased random walks, Levy flights, and fractal Brownian motion with long range memory (*H* = 0.9)(see Methods for implementations). We then analyzed the simulated data to determine if the feature set is sufficient to place each model in a distinct region of a joint feature space. Visualization of high-dimensional data sets, such as the Heteromotility feature set, presents a fundamental challenge. Here we employ *t*-Stochastic Neighbor Embedding (t-SNE), a visualization method that embeds high-dimensional data into a low-dimensional map, to generate 2D projections of our high dimensional feature space [45]. When Heteromotility feature space is visualized using t-SNE, these models of motion are clearly separated (Fig. 1A). Applying linear dimensionality reduction techniques such as principal component analysis (PCA)(Fig. 1B) and independent component analysis (ICA)(Fig. 1C) also provides some degree of cluster definition. Unsupervised hierarchical clustering (Ward’s method) [46] segregates the various models with high accuracy based on Heteromotility features (Fig. S2).

**Figure 1:**
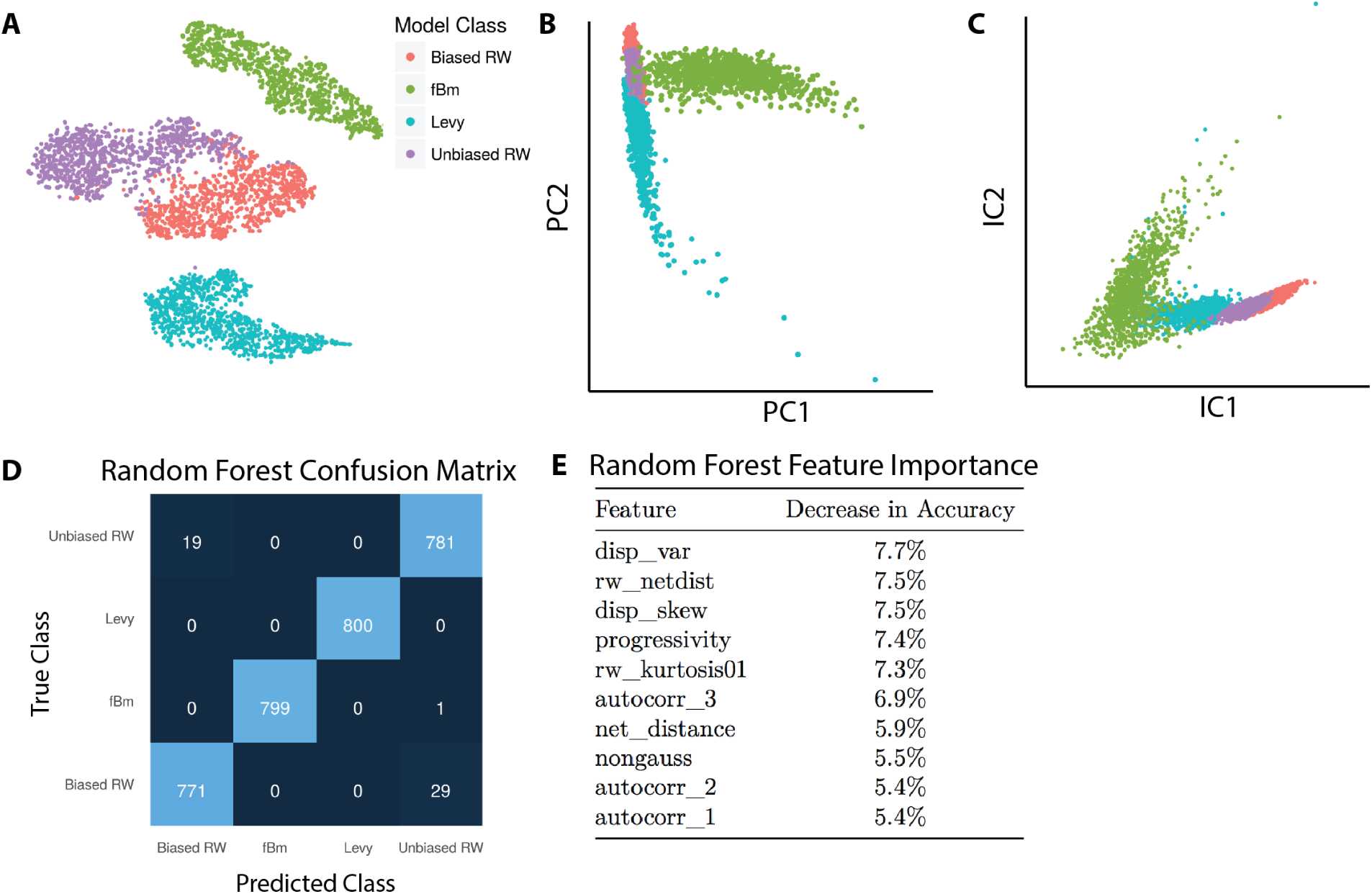
Simulated models of motion can be differentiated based on Heteromotility features. (A) t-SNE visualization of Heteromotility feature space determined for four simulated models, demonstrating clear separation of simulated data. (B) PCA visualization of motility feature space and (C) ICA visualization of motility feature space demonstrating that simulated data can be separated using simple dimensionality reduction. (D) The Confusion Matrix of a Random Forest classifier trained to distinguish the four simulated models of motion, mean accuracy of 98.4% (5-fold CV). (E) Features ranked by importance for Random Forest classification, where importance is determined as the decrease in accuracy when the feature is removed.

Applying a supervised Random Forest classifier [47], we are able to discriminate the four models of motion with 98.4% accuracy (5-fold cross validation score). We present the Random Forest classifier’s performance in a confusion matrix. A confusion matrix describes the errors made during classification in a matrix by listing the true classes as rows and the predicted classes as columns with values of the matrix cells representing the number of observations for each true class: predicted class pair (Fig. 1D) [48]. In this way, values along the matrix diagonal represent correct predictions, and values off the diagonal represent incorrect predictions. As seen in the matrix, there is little confusion between Levy flights, fBm, and the random walks, but some confusion between unbiased and biased random walks. The features important for effective classification can be determined by measuring the decrease in accuracy when each feature is removed. We find that while net distance contributed to two of the ten most important features for discrimination, the remaining eight features were metrics of the displacement distribution and autocorrelation, suggesting that Heteromotility’s additional features provide discriminative power beyond a simple speed/distance feature set (Fig. 1E). These results demonstrate robust detection of heterogenous motility phenotypes using a rich space of motion features. As a proof-of-concept, this analysis of simulated motion indicates that the Heteromotility feature set is sufficient to recognize different motility phenotypes where they exist.

### Wild-type and transformed MEFs occupy a shared region of motility state-space

Utilizing the Heteromotility software, we are able to analyze behavior in live cells and map this behavior into a cell state space. We initially sought to determine if this cell behavior state space was continuous, with cell states existing along a spectrum, or discrete, with a limited set of states cells could adopt. To answer this question, we first employed the mouse embryonic fibroblast culture system (MEF). Wild-type MEFs (WT MEFs) and MEFs transformed with *c*-*Myc* and *HRas*-*V12* overexpression constructs (MycRas MEFs) [49], which serve as a cancer model, were timelapse imaged for 8 hours to capture motility behavior. Images were segmented and tracked (see Methods), with cell centroid coordinates used as the cell location for tracking. By considering cell centroids, process extensions and retractions are considered as motile behaviors [26]. Analysis of cell paths with the Heteromotility software revealed that WT and MycRas MEFs differentially occupy different sections of a shared motility space, as visualized by t-SNE (Fig. 2A). Strikingly, MycRas MEFs occupy a sub-region of wild-type motility space, rather than a unique region.

**Figure 2:**
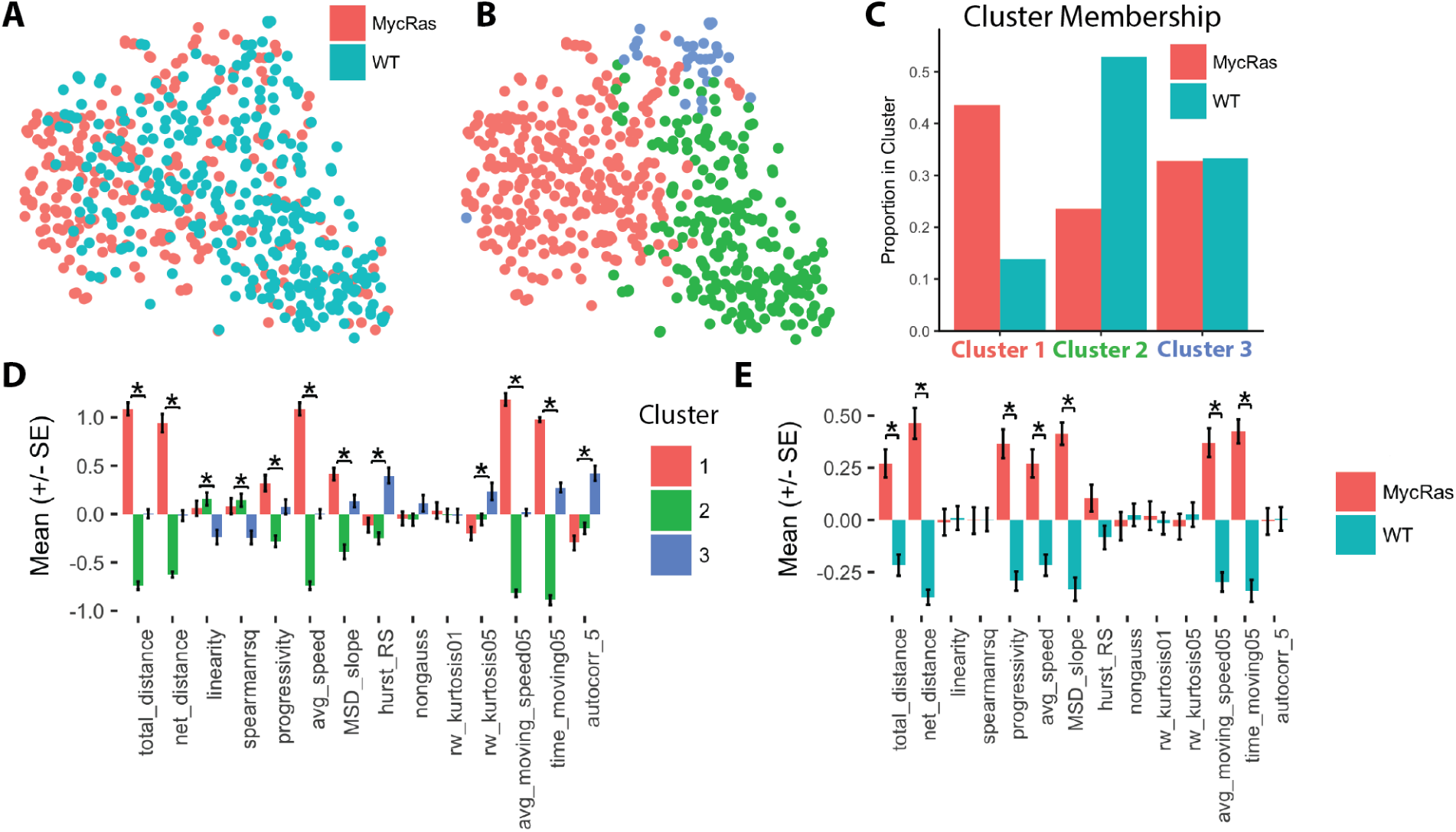
Wild-type and Myc/Ras transformed MEFs show differential occupancy within a shared motility state-space. (A) t-SNE visualization of wild-type (blue) and Myc/Ras transformed (red) MEFs in motility space. (B) Hierarchical clusters visualized with t-SNE (MANOVA, *p* < 0.001; Silhouette *S_i_* = 0.21). (C) Proportion of wild-type vs. transformed cells occupying each cluster. (D) Comparison of a subset of normalized feature values between clusters (^⋆^: Holm-Bonferroni corrected *p* < 0.01 by ANOVA) and (E) wild-type and transformed MEFs (^⋆^: Holm-Bonferroni corrected *p* < 0.01 by i-test). *n* > 250 cells per condition (pooled) from six independent experiments.

We applied hierarchical clustering (Ward’s method) [46] to the motility state space generated from pooled WT and MycRas MEFs to identify heterogenous motility phenotypes (Fig. 2B), considering a set of cluster validation indices to select the optimal number of clusters (See Supp. Methods). The Silhoutte value is a metric of cluster validity on the interval [-1, 1], taking into account the similarity of samples within a cluster and the difference between clusters [50]. Higher values indicate that samples within a cluster are similar and clusters are distinct. Notably, the cluster partition displays a positive Silhouette value, indicating an appropriate cluster structure. Feature mean values between clusters are also confirmed to be significantly different by multivariate analysis of variance (MANOVA) [51]. A complete set of cluster validation metrics is provided (Table S1). This result suggests that the state space is continuous, but can be decomposed into a set of overlapping states with characteristic behavioral phenotypes.

WT and MycRas MEFs are found to differentially occupy different clusters within the shared set, as may be expected from their initial distributions in the space (Fig. 2C). MycRas MEFs preferentially occupy Cluster 1, characterized by the highest average speed, proportion of time spent moving, and distance traveled, indicating a motile state. Conversely, WT MEFs preferentially occupy Cluster 2, characterized by the lowest average speed and time spent moving, and progressivity (net distance / total distance), indicating a less motile state (Fig. 2D). A small proportion of both MycRas and WT MEFs occupy Cluster 3, a rare population characterized by high kurtosis and progressivity, indicating a Levy flight like motile state that performs “jumping” motions. Observed visually, Cluster 2 cells are relatively immotile, Cluster 1 cells progress smoothly, and Cluster 3 cells exhibit erratic “jumping” motion (Supp. Movie 1).

MycRas and WT MEF cluster preferences are statistically significant by Pearson’s *χ*^2^ test of the *transformation state* × *cluster* contingency table (*p* < 0.0001). Considered as a population, the MycRas MEFs demonstrate higher progressivity, mean squared displacement, time moving, average speed, and self-similarity metrics than WT MEFs. This indicates that as a population they spend more time moving in a directed manner (Fig. 2E).

These results suggest that high motility and low motility states exist in both wild-type and transformed MEFs. Oncogenic transformation by *c*-*Myc* and *HRas*-*V12* may then be viewed as an input that leads a larger proportion of cells to adopt the high motility state, rather than introducing a novel phenotype unseen in WT cells. This observation is consistent with studies of motility behavior in the context of cancer, which have long suggested that increased motility is a phenotype of malignant cells [52,53]. Importantly, the motility behavior of a cancer cell population may be indicative of disease progression and outcomes [54]. We trained supervised classification models to predict if a cell was wild-type or transformed based on its motility behavior. Support vector machine (SVM) classifiers are a set of models that learn a decision boundary between classes. SVMs have shown efficacy in many problem domains [55]. Using an SVM classifier on the top 60% features selected by ANOVA F-value, we are able to classify individual cells as wild-type or MycRas transformed with ~75% accuracy based on motility features alone.

### Myoblasts display distinct motility states robust to perturbation

Our long-term goal is to understand the cell as a state machine, which entails not just identifying states, but also determining how various inputs from the extracellular environment may drive transitions between states. Mouse myogenic progenitor cells provide a well-characterized system in which chemical signals alter the behaviors of a mechanically-active cell type [56-59], and dynamic activation and lineage commitment processes can be manipulated [60,61]. In light of this long-term goal, we applied Heteromotility analysis to the myogenic system to ask how myogenic cells occupy state space.

Myoblasts are the transit amplifying progenitor of the skeletal muscle, produced as the daughters of muscle stem cells [60,61]. Myoblast motility has direct functional relevance, as muscle progenitors translocate along the muscle fiber to sites of injury during muscle regeneration [58,62] and transverse fibers during development [63]. Primary myoblasts were timelapse imaged for 8 hours, with and without stimulation by the growth factor FGF2. FGF2 is a known mitogen, inhibitor of differentiation, and possible chemoattractant in myogenic cells in culture [64-66]. *In vivo*, FGF2 is released during muscle injury [67,68], promoting expansion of the myogenic progenitor pool and possibly migration to sites of injury. As FGF2 is known to elicit different functional behaviors in myogenic cells, introducing this perturbation allows us to evaluate the robustness of myoblast motility states.

Applying hierarchical clustering, two motility clusters are detected (Fig. 3A). This partitioning scheme has a positive Silhouette value [50], significantly different cluster means by MANOVA, and optimizes cluster validation metrics (Table S1). Cluster 1 is a less motile state characterized by lower average moving speed, distance traveled, and a higher kurtosis. Conversely, Cluster 2 is characterized by a higher total distance, average speed, mean-squared displacement (MSD), and proportion of time spent moving. These characteristics indicate that myoblasts occupy a two state system with a consistently motile state (Cluster 2), and a less motile state that exhibits more Levy-flight like displacement behavior (Cluster 1). Observed in videos, cells in Cluster 1 are less motile, and display less directed motion than cells in Cluster 2 (Supp. Movie 2).

**Figure 3:**
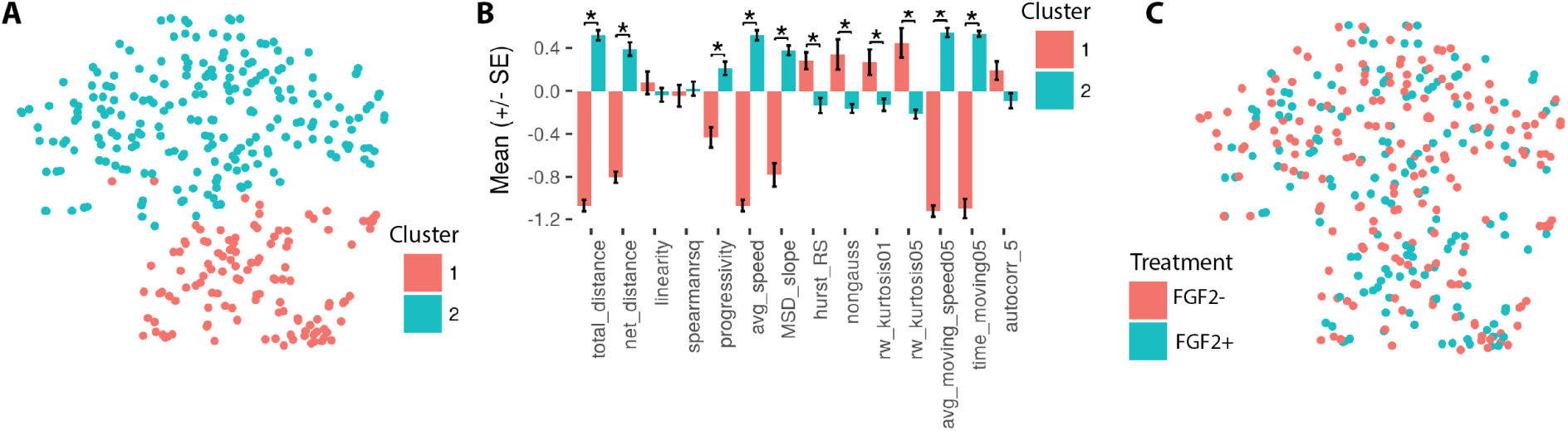
Myoblasts display distinct motility states, shared by both FGF2+ and FGF- conditions. (A) t-SNE visualization of hierarchical clusters in myoblast motility space (MANOVA, *p* < 0.001; Silhouette *S_i_* = 0.30). (B) Normalized feature means for motility state clusters (^∗^: *p* < 0.05, Holm-Bonferroni corrected *t*-test). (C) t-SNE visualization of FGF2 treated (blue) and untreated (red) myoblasts in motility space. *n* ~ 150 cells per condition, taken from two separate animals, with total *n* = 338 cells.

Both FGF2 stimulated (FGF2+) and unstimulated (FGF2-) cells co-occupy both states, with no notable preference in state space induced by either condition (Fig. 3C). This is confirmed quantitatively as a lack of preferential cluster occupancy between FGF2 treated and untreated cells (*χ*^2^ test *p* > 0.05 of *FGF2*-*treatment* × *cluster* contingency table). This result suggests that FGF2 does not induce a notable effect on myoblast motility under these conditions.

### Muscle stem cells display motility states reflecting activation

Until this point, the cells analyzed have not been undergoing any dramatic phenotypic transitions. In contrast, stem cells are specifically designed to undergo dynamic activation and differentiation processes that radically reshape cell phenotype on an hours to days long timescale. Would it be possible to observe such phenotypic transitions through the lens of cell behavior changes? We apply Heteromotility analysis to the muscle stem cell (MuSC) system during activation from quiescence in an attempt to observe dynamic transitions in cell behavior. MuSCs undergo an exit from quiescence in cell culture conditions, entering an activated state over a roughly 48 hour window, providing a model of a dynamic process where behavior state transitions would be expected [60,69]. Heterogeneity within the MuSC population is well appreciated [11], suggesting that motility behavior may also be heterogenous during activation. Motility behavior is also relevant to MuSC function due to the physiological motility behavior of muscle progenitors during regeneration, as noted above.

Primary MuSCs were isolated from limb muscles by FACS (PI^-^/CD31^-^/CD45^-^/Sca1^-^/VCAM^+^/*α*7-integrin^+^) [70] and seeded on sarcoma-derived ECM coated well plates. After 24 hours in culture, MuSCs were timelapse imaged for 8 hours in DIC. At this stage, MuSCs have begun to activate (*MyoD^+^*), but are not yet committed to differentiation (*MyoG^+^*) [69]. Visualizing hierarchical clusters (Ward’s method) of MuSC motility features with t-SNE, it is apparent that multiple motility subpopulations are present (Fig. 4A). We identify four distinct clusters with notably different phenotypes. This partitioning scheme has a positive Silhouette value [50], significantly different cluster means by MANOVA, and optimizes cluster validation metrics, as above (Table S1). As in myoblasts, the clusters appear to separate based on differences in total distance, average speed, and time moving. Additionally, clusters segregate based on the linearity and progressivity of motion.

Cluster 1 is characterized by the lowest total distance, average speed, time moving, progressivity, and MSD. Observed visually, cells in Cluster 1 are immotile and appear morphologically rounded, lacking any filopodia characteristic of myogenic activation and motility (Supp. Movie 3). Clusters 2, 3, and 4 are characterized by increasing measures of total distance, average speed, and time spent moving. Cluster 4 exhibits the highest of these metrics and high kurtosis, suggesting a highly motile Levy-flight like state. Cluster 2 exhibits higher progressivity, linearity, and MSD, indicating a more directed motility state (Fig. 4D)(*p* < 0.001 by ANOVA for all features displayed).

**Figure 4:**
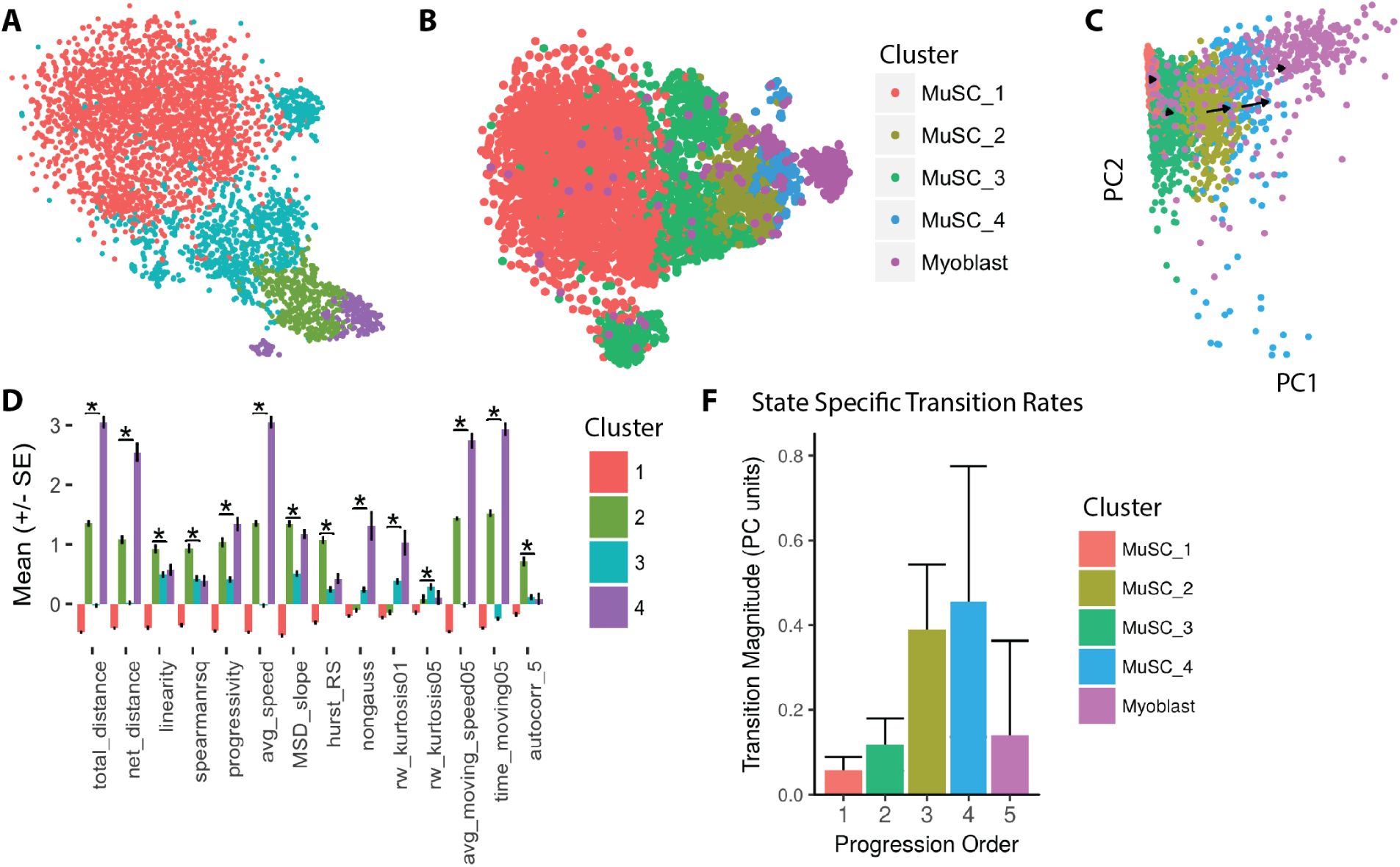
MuSC motility states reflect progressive myogenic activation. (A) t-SNE visualization of distinct motility states, as detected by hierarchical clustering (colors)(MANOVA, *p* < 0.001; Silhouette *S_i_* = 0.18). (B) t-SNE visualization of MuSC motility states and myoblasts in a shared space suggesting progressive myogenic motility states. (C) Average transition rate vectors for MuSC motility states (arrows, scaled for presentation) demonstrating transitions toward the myoblast motility phenotype over time in a shared PCA space. (D) subset of normalized feature means for each motility state (^∗^: p < 0.05, Holm-Bonferroni corrected ANOVA). (F) The magnitude (+/- SEM) of the mean transition vector for all cells in a given state. All data represent analysis of *n* > 1800 cells per condition pooled from three animals, with total *n* = 6557 cells.

MuSCs incubated with FGF2 display a similar distribution in motility space compared to untreated cells (Fig. S3). Quantitatively, FGF2 MuSCs display no motility state preference relative to untreated cells (Fig. S3)(*p* > 0.05, *χ*^2^ test of *FGF2*-*treatment × cluster* contingency table). On a population level, FGF2 does not appear to significantly alter any of the motility features calculated by our software, suggesting that it may not play a role in regulating motility in these conditions at this stage of myogenic activation.

Visualized in a shared t-SNE space with myoblasts, the order of MuSC motility states suggests a progressive motion toward the myoblast state space as total distance, time moving, and speed metrics increase (Fig. 4B). This is confirmed in a linear state space using principal component analysis (PCA), in which MuSC states and myoblasts segregate primarily along the first principal component (PC1)(Fig. 4C). The top 20 features contributing to PC1 in this shared space are metrics of moving speed and time spent moving. This result indicates that MuSC motility states reflect progressive states of activation toward the eventual myoblast motility state space. We explored this possibility with pseudotime analysis, which attempts to reconstruct a dynamic cellular process from high-dimensional single cell data under the assumption that the process is ergodic and cells move in a continuous manner through feature space over time [18,71]. Analytically, this is most commonly achieved by fitting a minimum spanning tree through the data in reduced dimensional space, interpreting the tree’s longest axis as a temporal axis. Pseudotime analysis in the MuSC motility state space reflects the view that MuSC states are ordered in a progressive series (Fig. S3).

To further confirm that the MuSC motility states reflect states of activation, we quantified state transition rates and directions for cells in each MuSC state and myoblasts. Cell paths were divided into three equal length tracks ( *τ* = 20 frames) and Heteromotility features were extracted for each of these subpaths. The state of a cell during each time interval *τ* was defined in two dimensions as the cell’s location along the first two principal components of a shared MuSC and myoblast PCA space. Transitions for each cell were calculated as the vector between sequential 2D state locations. The mean transition vector for a given cluster was calculated as the mean of all transitions made by all cells assigned to that cluster.

Visualizing transition rates as vectors originating from each cluster centroid in PCA space, it is evident that cells in each MuSC state progress toward the next state in the sequence over time (Fig. 4C). The myoblast motility states are present at the end of the sequence, suggesting that MuSC states represent different stages of myogenic activation. Notably, the transition rates increase as a function of state progression, suggesting the kinetics of activation are non-linear (Fig. 4F). In this way, we provide quantitative measurements of transitions between states of myogenic activation in single cells for the first time.

### MuSC motility states are more dynamic than MEF and myoblast motility states

We next sought to investigate the dynamics of our motility state systems. A key benefit of using behavior features to define states is that a single cell can be subjected to a state assay at multiple time points, allowing state transitions to be detected. How long does a cell reside in a particular state? Can every state transition to every other state, or are transitions restricted to exist between particular states, as would be the case in a finite state automaton? The answers to such questions would help clarify the computational logic underlying cell behavior.

To visualize and quantify the dynamics of the motility state systems, we applied coarse-grained probability flux analysis (cgPFA), as implemented by Battle *et. al.* [72] (Supp. Methods). In this method, the first (PC1) and second (PC2) principal components are segmented into *n* bins, and we define each unique combination of bins on PC1 and PC2 as a unique state. To define cell state at multiple points using motility features, cell paths are segmented into *τ* length subpaths and motility features are extracted from each subpath. Cell state is defined for each subpath based on its position in coarse-grained PCA space, where each binned coordinate is treated as a state. Cell states are compared from one subpath to the next to quantify the dynamics of the motility state system (see Supp. Methods).

As a validation of our implementation, we performed cgPFA on simulated cell paths that varied their model of motility on a defined interval (*τ* = 20 time points) and compared them to invariant simulations, using subpaths of the same length as the variable model’s states (*τ* = 20 time points). For each bin, a state transition rate is calculated as the vector mean of transitions from that state bin into neighboring states in the von Neumann neighborhood. These transition rates are visualized as arrows atop each state bin. Arrow direction represents the direction of transition rate and arrow length represents the rate magnitude. State bins with high transition consensus therefore display longer arrows, indicating that most cells transition in the same direction. As a measure of state stability, the divergence of the vector field is displayed as a heatmap. States with negative divergence may be considered metastable, with more cells entering than exiting. A Levy flight model transitioning to a random walk displays a highly directed set of state transitions (Fig. 5A), as compared to a simple random walk which displays minimal directionality (Fig. 5B). Quantifying the directionality of the transition vector field as the weighted mean magnitude of the transition rate vectors, the variable model displays higher directionality than the invariant model as expected (Fig. 5C).

**Figure 5:**
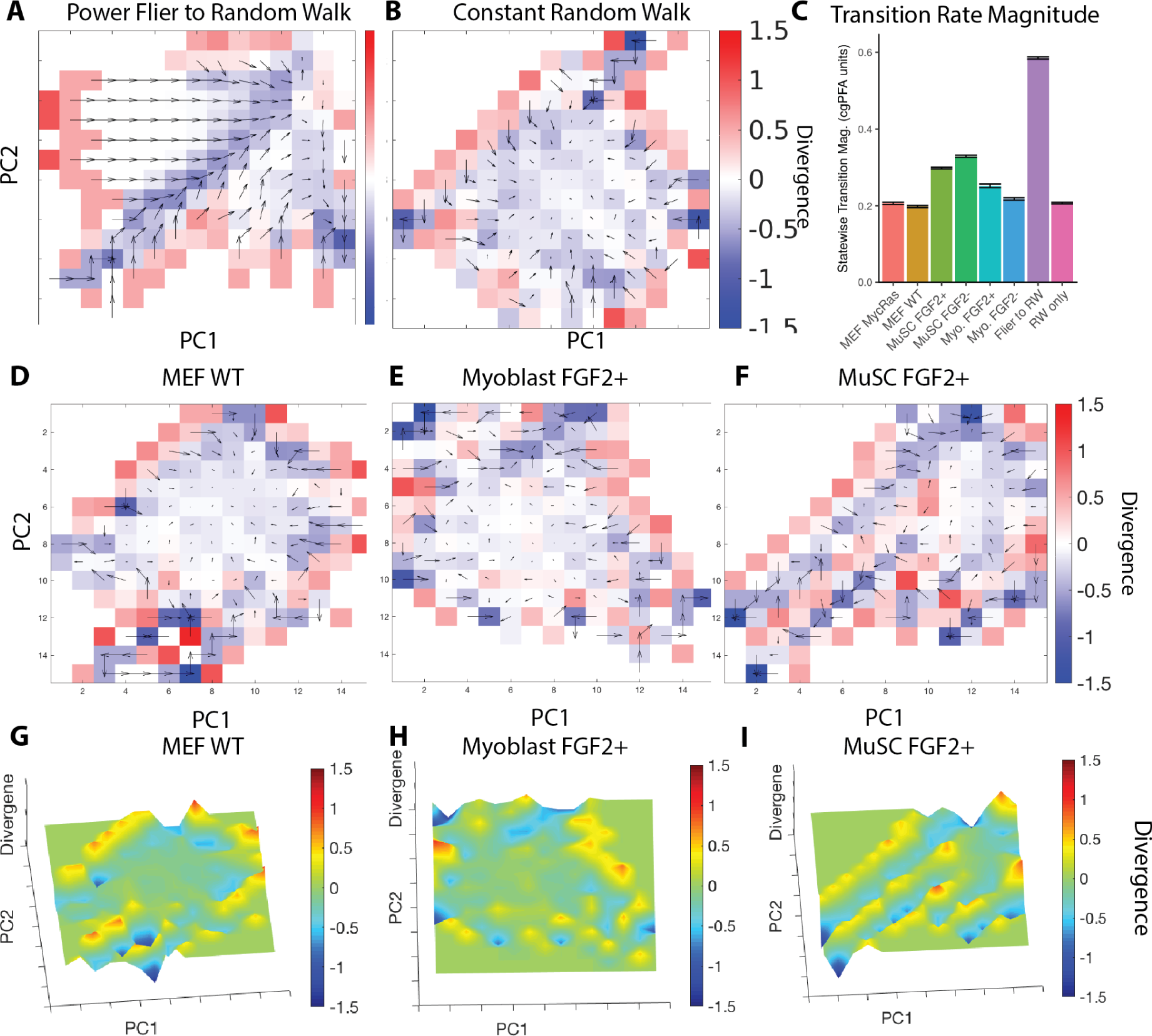
coarse-grained probability flux analysis (cgPFA) of (A) Levy flier to random walk simulations and (B) Random walk invariant simulations. cgPFA of (D) WT MEF, (E) Myoblast FGF2+, and (F) MuSC FGF2+ motility states with subpaths of length *τ* = 20 time points (130 minutes). Each unique combination of bins between PC1 and PC2 is considered as a unique state. Arrows represent transition rate vectors, calculated for each state bin as the vector mean of transitions into the neighboring states in the von Neumann neighborhood. Arrow direction represents the direction of these transition rate vectors, and arrow length represents transition rate vector magnitude. Underlying colors represent the vector divergence from that state as a metric of state stability. Positive divergence indicates cells are more likely to leave a state, while negative divergence indicates cells are more likely to enter a state. (C) Net flow of coarse-grained motility space, estimated as the magnitude of the vector sum of transition rate constants. 3D representations of (G) MEF WT, (H) Myoblast FGF2+, and (I) MuSC FGF2+ motility state divergence.

Applying cgPFA to our biological systems with subpaths of length *τ* = 20 time points (130 minutes), both WT MEF and MycRas MEF systems display no obvious state flux and directionality on par with the invariant random walk simulation (Fig. 5D, S4). Topographically, MEF systems display a ‘basin’ of metastable states. This region has near zero divergence and low transition rates. States on the outer periphery of this metastable region have higher divergence and transition rates, indicating that these states are less stable (Fig. 5D, S4). Visualizing state space divergence in three dimensions, the metastable states appear as a central valley, while the unstable states appear as peaks (Fig. 5G). Myoblast motility state systems appear similar to the MEF systems by simple observation. Transition directionality is only slightly higher than the invariant simulation (Fig. 5C), and the topology appears to be dominated by a metastable basin at the center, surrounded by unstable states on the edge of this basin. Topology and transition directionality are comparable in FGF2 treated and untreated conditions (Fig. 5E, S4).

MuSC motility state systems appear more dynamic than MEF and myoblast systems, as indicated by higher transition directionality (Fig. 5C). This transition directionality is readily apparent when viewing the transition vector fields, with most transition rates leading toward two metastable ‘valleys’ and a metastable basin on the edge of state space (Fig. 5F, S4). Visualized in three dimensions, these metastable regions are clearly visible as valleys in state space bordered by unstable ‘ridges’. To determine how long MuSCs occupy a given state, we characterized the dwell times of each occupied state in the course-grained PCA space by exponential decay curve fitting, providing a time constant for each state. In these course-grained state spaces, dwell times appear exponentially distributed with time constants ranging from *τ*_*TC*_ ≈ 0.6 to *τ*_*TC*_ ≈ 1.3 (Fig. S6). The apparent exponential distribution of dwell times suggest that the state transition process is Markovian on the timescales we observe. Time constants in this range indicate that the majority of cells transition between course-grained states within a single discrete time unit. Time constants are positively correlated with the number of cells occupying a state, supporting the notion that states of high occupancy are metastable and therefore characterized by longer dwell times (Fig. S6). Topology, dwell times, and transition directionality are comparable between FGF2 treated and untreated conditions (Fig. 5I, S4, S6). These results suggest that the MuSC motility state system is more dynamic than the MEF or myoblast systems, consistent with the dynamic MuSC activation process, and displays Markovian transition dynamics.

### MuSC motility states break detailed balance

A system with a discrete set of states at equilibrium should obey the law of detailed balance, such that each individual state transition *A* → *B* occurs at the same rate as the reverse transition *B* → *A.* Systems that breaks detailed balance transition between states in a directed manner, such that future behavior can be predicted by current state. Biological systems frequently break equilibrium when undergoing directed processes, but confirming that detailed balance is broken in a given scenario can prove challenging [72]. A system in detailed balance would be expected to exhibit equal and opposite transition vectors with no observable pattern of transitions. In our MuSC systems, visualizing transition rates by PFA suggests that detailed balance may be broken, as shown by the directed nature of transition rate vectors (Fig. 5, S4).

To confirm statistically that detailed balance is broken in these systems, we defined *N*-dimensional state spaces based on the first *N* coarse-grained principal components and performed PFA in these spaces (ND-cgPFA)(see Methods). A 2*N*-dimensional matrix is generated representing every possible set of state transitions in a given *N* -dimensional state space. For example, a 1 dimensional state space is represented as a 2 dimensional matrix, each dimension coarse-grained into *k* bins (rows, columns). One dimension of this 2D space represents a cell’s initial state position at one time point, and the other represents the destination state position at the next time point (as in Fig. 6A, B). This pattern is repeated for the construction of higher dimensional spaces. For instance, a 2D state space is represented as a 4D matrix, where the *first two* dimensions describe initial state, and the *next two* dimensions describe final state. The number of cells that exhibit each transition are recorded as the value of the corresponding position in the matrix (see Methods for further description).

**Figure 6:**
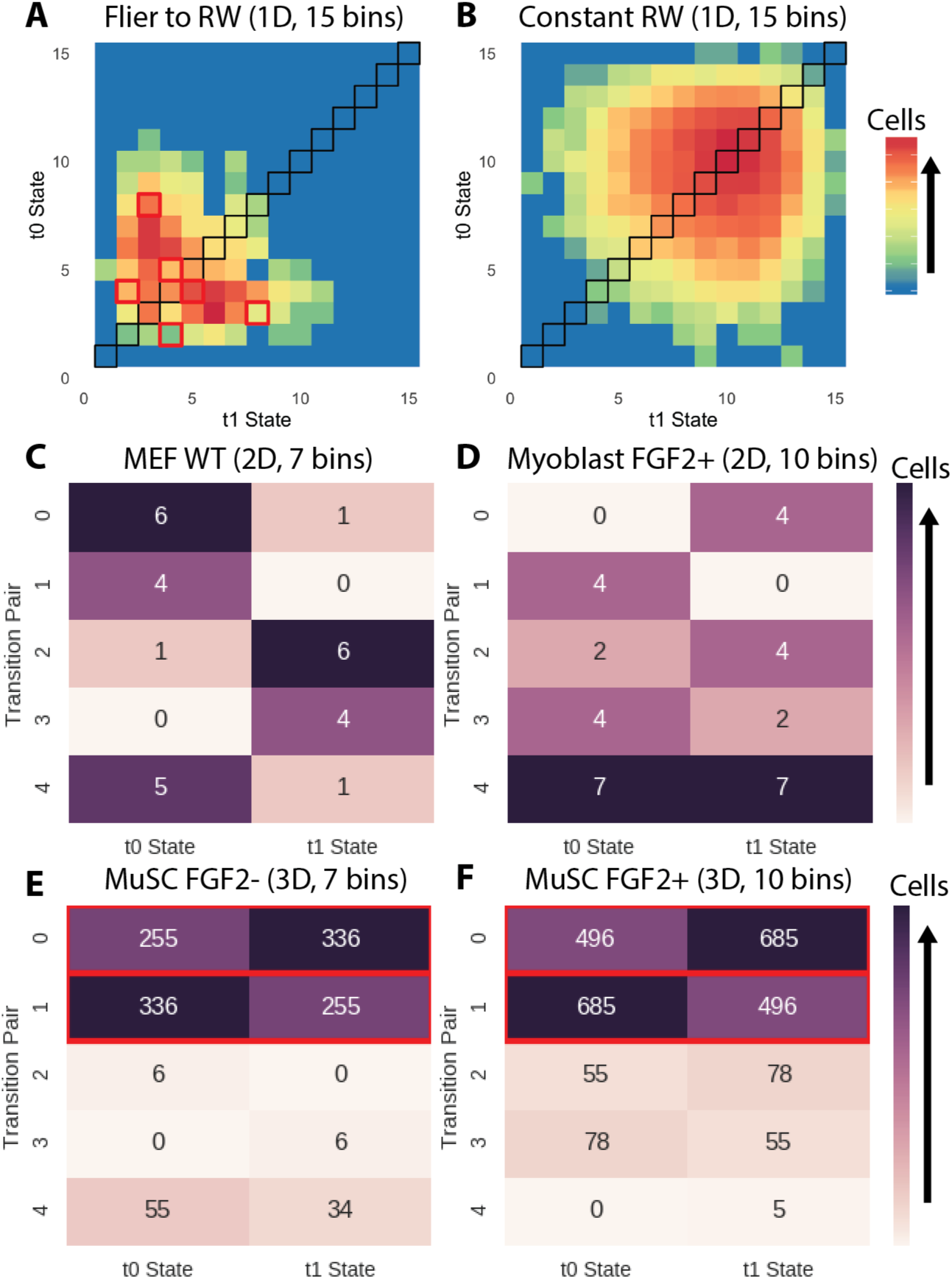
One-dimensional coarse-grained PFA of (A) Simulated Levy flier transition to a random walk, (B) simulated random walk. Transition pairs from *N*-dimensional cgPFA displayed as heatmaps for (C) WT MEFs (*n* = 312), (D) Myoblasts (FGF2+) (*n* = 150), (E) MuSCs (FGF2-) (*n* = 1838), and (F) MuSCs (FGF2+) (*n* = 2500). Heatmaps show the five most unbalanced transitions in a system, with colors and numerical insets indicating the number of cells that transitioned from a given state at *t*_0_ to a given state at *t*_1_. Significantly unbalanced transitions are outlined in red (*p* < 0.05, Holm-Bonferroni corrected binomial test). A system in detailed balance would display no unbalanced transitions.

A system in detailed balance would be expected to display symmetry for each pairwise set of transitions. To determine if each of our systems was in detailed balance, we performed ND-cgPFA at several levels of dimensional (*N* ∈ {1, 2, 3, 4}) and coarse-grained (*bins k* {2, 3, 5, 7, 10, 15, 20}) resolution. At each level of resolution, we test each of the *k*^2*N*^ possible pairwise transitions for balance by the binomial test (*H*_0_: *p* = 0.50).

To validate this method, we performed cgPFA on simulated cell paths generated using variable and invariant models of motion. A Levy flier that transitions to a random walk shows clearly unbalanced pairwise transitions (Fig. 6A) at one dimension of resolution, while an invariant random walk displays symmetry about the diagonal and balanced pairwise transitions (Fig. 6B). A simple binomial test for pairwise transitions finds multiple unbalanced transition pairs in the variant model, but not the invariant model (Fig. 6A, B).

A-dimensional matrices cannot be visualized in their totality as in the 1D case. As noted above, we tested all *k*^2*N*^ possible transitions for balance at multiple dimensional and binning resolutions for each system. Here, the five transition pairs that are most unbalanced out of the possible set of *k*^2*N*^ transitions (by *p* value of the binomial test) in a given system and state space are presented as a 5-by-2 heatmap, where columns represent the initial and final states in a transition pair. We present the dimensional and coarse-grained resolution revealing the most asymmetry for each system, as noted above each heatmap. Visualizing ND-cgPFA pairwise transitions for MEF (Fig. 6C), myoblast (Fig. 6D), and MuSC state systems (Fig. 6E, F) demonstrates that transitions with some degree of unbalance exist in each of the systems. By the binomial test (Holm-Bonferroni corrected), only the MuSC systems (FGF2+ and FGF2-) display significantly unbalanced transitions in any of the course-graining schemes tested, confirming that detailed balance is broken. Our ND-PFA test is biased toward Type II error (false negatives) rather than Type I error (false positives) for multiple reasons, providing further confidence that the detailed balance breaking we identify is valid. It is therefore possible that detailed balance is also broken in the MEF and myoblast systems and our tools are simply not sufficient to detect this asymmetry.

As additional confirmation of MuSC motility state dynamism, we performed this symmetry breaking analysis using hierarchical clustering to define cell state over the whole motility feature space, rather than coarse-grained location along the principal components (Fig. S5). As with ND-cgPFA, our positive control variable model breaks detailed balance by the binomial test, while our negative control invariant models do not. MuSC systems again demonstrate greater pairwise asymmetry than MEF or myoblast systems in this test (Fig. S5).

An important question is raised when detailed balance is broken. Is the system stationary, with the same number of cells in each state over time, or non-stationary? We evaluated the stationary nature of the MuSC systems by forming a contingency table comparing state occupancy between time points for each set of length *τ* = 20 subpath the dataset and find a significant difference (*p* < 0.05, *χ*^2^ test) for each scale where detailed balance breaking is detected. These results collectively demonstrate the more dynamic nature of the MuSC system and demonstrate that the MuSC motility state system is non-stationary and breaks detailed balance.

## Discussion

Cell behavior phenotypes represent an under-exploited opportunity to explore the heterogeneity of cell systems. In contrast to existing methods based on destructive molecular assays, cell behaviors such as motility can be tracked in a single cell over time, allowing for measurement of phenotypic state transitions. Our Heteromotility analysis software provides access to one portion of this rich feature space for analysis. Applying analysis techniques common in other single cell assays, we demonstrate that Heteromotility features are sufficient to distinguish different motility phenotypes and provide novel insight in multiple biological contexts. In addition to detecting heterogeneity within populations, Heteromotility analysis was also useful to quantitatively describe perturbations on the population level.

In our three biological systems, overlapping but characteristic motility states are present in a continuous state space. Notably, a shared set of motility states is conserved between WT and MycRas MEFs, despite the dramatic perturbation of neoplastic transformation. A similar phenomenon of conserved phenotypic states has been described for cell shapes. Despite a large number of perturbations, multiple cell systems displayed a fairly limited set of cell morphology states, with perturbation merely altering the prevalence of these states [4,73]. These results suggest that cell systems may have a discrete set of phenotypic states despite a much greater diversity in molecular organization, with perturbations acting largely to alter the distribution of cells within these states, rather than elicit novel behavior.

Although these state definitions are robust to perturbation in our conditions, the distribution of cells among these states appears to be dynamic. This is demonstrated by the state preferences of WT and MycRas transformed MEFs. Considering the robustness of motility state definitions, state transitions may act as a mechanism of population level phenotype change. This is not necessarily in opposition to a model of motility regulation in which cell phenotypes shift within a given state. Harkening to the subsumption architectures of robotic control systems [74], a higher level state determinant, such as oncogene expression, may be viewed as inducing a preference for the selection of behavioral states, and substates within those states, by a more direct effector mechanism, in this case the cell motility machinery. The mechanisms of state transitions and substate preference may work in synchrony to specify population level phenotypes.

Within the MEF systems transformed cells preferentially occupy the more motile state, suggesting that neoplastic transformation shifts the distribution of cells within the motility state system. This state preference is strong enough to allow for better-than-chance machine learning classification of WT and MycRas MEFs based solely on motility features. Training machine learning classifiers on quantitative motility features and ground truth patient outcomes may potentially serve as an additional cancer diagnostic metric.

Most single cell analysis methods currently used to investigate heterogeneity are not capable of assaying a single cell across a time course, but rather provide a detailed snapshot of a cell at a single time point. Real-time observation of individual cells may elucidate novel properties of a heterogenous cell system. Here, we combine the real-time nature of cell behavior observations with Heteromotility analysis to investigate the dynamics of motility states.

In the context of MuSCs undergoing myogenic activation, we find a set of motility behavior states progressively more similar to myoblast motility behavior. Quantifying state transitions within each of these states, we observe that cells within each state transition toward the next state in the series over time. State transition rates increase as a function of state progression, suggesting that MuSC activation is not simply a linear process. This observation may have implications for the study of MuSC heterogeneity. If the rate of phenotypic change in MuSCs is non-linear during activation, then heterogeneity in activation kinetics between cells will be exaggerated during a ‘critical period,’ where the most rapidly activating cells are not only more activated at that moment, but are moving toward a fully activated state more rapidly than their less activated counterparts. This is to our knowledge the first quantification of single cell transition rates between quiescent MuSC and myoblast phenotypes during myogenic activation. It follows that similar analysis of cell behavior may elucidate transition dynamics in other contexts of cell biology.

A state system at equilibrium displays detailed balance, in which pairwise transitions between all states are equal. We check for the presence of detailed balance in our motility state systems as a method of quantitatively discerning if cells are transitioning through state space in a predictable manner. The MuSC motility state system breaks detailed balance, while the MEF and myoblast systems do not. This may reflect underlying properties of each system, as MuSCs are undergoing a dynamic activation process on the timescale of imaging, while MEFs and myoblasts are not. Broken detailed balance in the MuSC system indicates that the current MuSC state provides predictive information about the cell’s future state, as cell’s transit state space in a predictable way. This supports our observation of progressive activation states discussed above. Analysis of detailed balance breaking by real-time cell state inference from cell behavior may be a useful approach to detect dynamic biological processes and predictable phenotypic patterns.

## Conclusions

Our Heteromotility software provides a means of defining cell states and quantifying transitions between them by quantitative analysis of one aspect of cell behavior. We demonstrate the analysis is capable of identifying unique motility states in simulations and three biological systems. In our biological systems, these states appear to have robust definitions when perturbed, but each cell’s state identify appears dynamic. Real-time, quantitative assays of single cells such as Heteromotility analysis may reveal the dynamics of heterogenous cell systems, which cannot be accomplished by terminal molecular assays. We demonstrate this approach by showing that MuSCs transition through a progressive set of motility states toward the myoblast motility state in a predictable manner, breaking detailed balance.

## Acknowledgements

This work was funded by NSF grant MCB-1515456 to W.F.M., NIH grants AR060868 and AR061002 to A.S.B., and a PhRMA Foundation fellowship to J.C.K. This material is based upon work supported by the National Science Foundation Graduate Research Fellowship under Grant No. 1650113 to J.C.K. J.C.K, A.C., and W.F.M. are members of the NSF Center for Cellular Construction, NSF Grant No. 1548297. The authors would like to thank Morgan L. Truitt and Davide Ruggero for generously providing transformed MEF cells.

## Materials and Methods

### Animals

All mice were housed at the University of California San Francisco following UCSF Institutional Use and Care of Animals Committee guidelines. Adult male C57Bl/6 mice (2-4 m.o.) were used for myogenic cell experiments. Adult female C57Bl/6 mice (2-3 m.o.) were used as mothers to derive MEFs from E13.5 embryos [75].

### Cell Isolation and Culture

Mononuclear cells were isolated from limb muscles of adult mice as described [76]. Myoblasts were isolated by negative-selection against *CD31*, *CD45*, and *Sca1* using the EasySep Endothelial Selection kit and subsequent 5 minute preplating. Myoblasts were maintained in myogenic growth media (F10, 20% FBS, and [5 ng/mL] FGF2) on sarcoma-derived ECM coated dishes. All myoblasts used in experiments were passage 3-4. Wild-type muscle stem cells were isolated by FACS (PI-/CD31-/CD45-/Sca1-/VCAM+/*α*7-integrin+) and cultured as described [70]. Wild-type MEFs were isolated from E13.5 embryos as described [75]. Transformed MEFs were generated as described [49], generously donated by the authors. All MEFs were maintained in DMEM, 10% FBS, 1% Penicillin / Streptomycin.

### Timelapse Motility Imaging

All images were performed in a temperature and CO2 controlled unit at 37^O^C and 5% CO_2_. All experiments were performed with a 20X air objective, capturing images with a Hamamatsu C11440-22CU camera (pixel size 6.5 × 6.5 *μm*) using a Nikon Ti with automated XY stage. MuSCs were seeded at 500-1000 cells/well in 20 wells of an optically clear tissue culture treated, plastic bottomed 96 well plate coated with sarcoma-derived ECM (Sigma) immediately after FACS sorting. After 23 hours to allow for adaptation to culture, media was exchanged for the relevant experimental medium and MuSCs were incubated for 1 hour to adapt. MuSCs were imaged at 20X in DIC for 10 hours with a temporal resolution of 6.5 minutes / frame. Myoblasts were passaged and plated 24 hours prior to imaging at 300-500 cells/well. Media exchanges were performed 1 hr prior to imaging, in the same manner as MuSCs. Myoblast imaging was performed in the same manner as MuSCs. MEFs were similarly plated 16-20 hours prior to imaging at the same density and imaged in the same manner. For FGF2 perturbation experiments, 10 wells of an optically clear plastic bottomed 96 well plate were imaged in growth media with [5 ng/mL] FGF2, and the remaining 10 were imaged in growth media with [0 ng/mL] FGF2. Additional details are available in the Supplemental Methods.

### Heteromotility Analysis

DIC images were segmented using custom segmentation algorithms, optimized for each of the systems we investigated. Tracking was performed with a modified version of uTrack [77]. Code is available on the Heteromotility GitHub page. All code for the Heteromotility tool is available on the Heteromotility Github page (https://github.com/jacobkimmel/heteromotility) and in the Python Package Index (PyPI). Detailed descriptions of the algorithms are provided in the Supplemental Methods.

